# Strain dependent differences in coordination of yeast signaling networks

**DOI:** 10.1101/2022.06.09.495559

**Authors:** Taylor D. Scott, Ping Xu, Megan N. McClean

**Affiliations:** Department of Biomedical Engineering, University of Wisconsin-Madison, Madison, WI, USA; Lewis-Sigler Institute for Integrative Biology, Princeton University, Princeton, NJ, USA; University of Wisconsin Carbone Cancer Center, University of Wisconsin School of Medicine and Public Health, Madison, WI, USA

## Abstract

The yeast mitogen activated protein kinase pathways serve as a model system for understanding how network interactions affect the way in which cells coordinate the response to multiple signals. We have quantitatively compared two yeast strain backgrounds YPH499 and Σ1278b (both of which have previously been used to study these pathways) and found several important differences in how they coordinate the interaction between the high osmolarity glycerol (HOG) and mating pathways. In the Σ1278b background, in response to simultaneous stimulus, mating pathway activation is dampened and delayed in a dose dependent manner. In the YPH499 background, only dampening is dose dependent. Further, leakage from the HOG pathway into the mating pathway (crosstalk) occurs during osmostress alone in the Σ1278b background only. The mitogen activated protein kinase Hog1p suppresses crosstalk late in an induction time course in both strains but does not affect the early crosstalk seen in the Σ1278b background. Finally, the kinase Rck2p plays a greater role suppressing late crosstalk in the Σ1278b background than in the YPH499 background. Our results demonstrate that comparisons between laboratory yeast strains provide an important resource for understanding how signaling network interactions are tuned by genetic variation without significant alteration to network structure.

## Introduction

Cells exist in dynamic changing environments and must coordinate their respond to multiple stimuli in order to survive and thrive. The cellular response to a stimulus may be organized into a series of tightly regulated reactions which together form a signaling network. The mitogen activated protein kinase (MAPK) network architecture is a common example of a signaling network and is conserved among eukaryotes [1]. The core structure consists of a three-step kinase cascade, in which a MAP kinase kinase kinase (MAP3K) becomes activated by diverse upstream mechanisms. Once activated, it phosphorylates and activates a MAP kinase kinase (MAP2K), which phosphorylates and activates the MAPK. Once activated, the MAPK phosphorylates diverse targets throughout the cell in order to effect the cellular response. Multiple MAPK networks may exist in a cell, each controlling the response to one or more stimuli, and these networks may share kinases. MAPK networks play an important role in many fields of study, including disease etiology and drug development [2], developmental biology [3], and *in silico* control of cellular processes[4].

The budding yeast (*Saccharomyces cerevisiae)* MAPK networks are an attractive model system for studying signal flow through MAPK networks because their stimuli are well known, and they have a relatively simple structure. Two pathways of interest are the high osmolarity glycerol (HOG) pathway, which controls the response to high osmotic stress, and the mating/FG pathway, which is activated by pheromone and nutrient starvation (Figure 1). The HOG pathway consists of two parallel activation branches, known as the SLN1 branch (containing the MAP3Ks Ssk2p and Ssk22p) [5,6] and the SHO1 branch (containing the MAP3K Ste11p) [6,7], which converge on the MAP2K Pbs2p. Pbs2p activates the sole MAPK, Hog1p [8]. Activation of the Hog1p can be measured via the transcriptional reporter *STL1*, which is strongly activated by HOG pathway activity in a Hog1p-dependent manner [9,10]. The mating/FG pathways share a single activation branch [11] in which the MAP3K Ste11p is activated by the Cdc42-Ste20 complex [12–16] before phosphorylating and activating the MAP2K Ste7p [17,18], which activates the MAPKs Fus3p and Kss1p [19–22]. While Fus3p and Kss1p are traditionally thought of as the MAPKs for the mating and FG pathways respectively, they are partially redundant in mating [23] and are both activated in response to pheromone [24–27]. *FUS1* is a common transcriptional reporter of mating/FG MAPK activity, and it is induced by both phosphorylated Fus3p and Kss1p [28].

**Figure 1:**
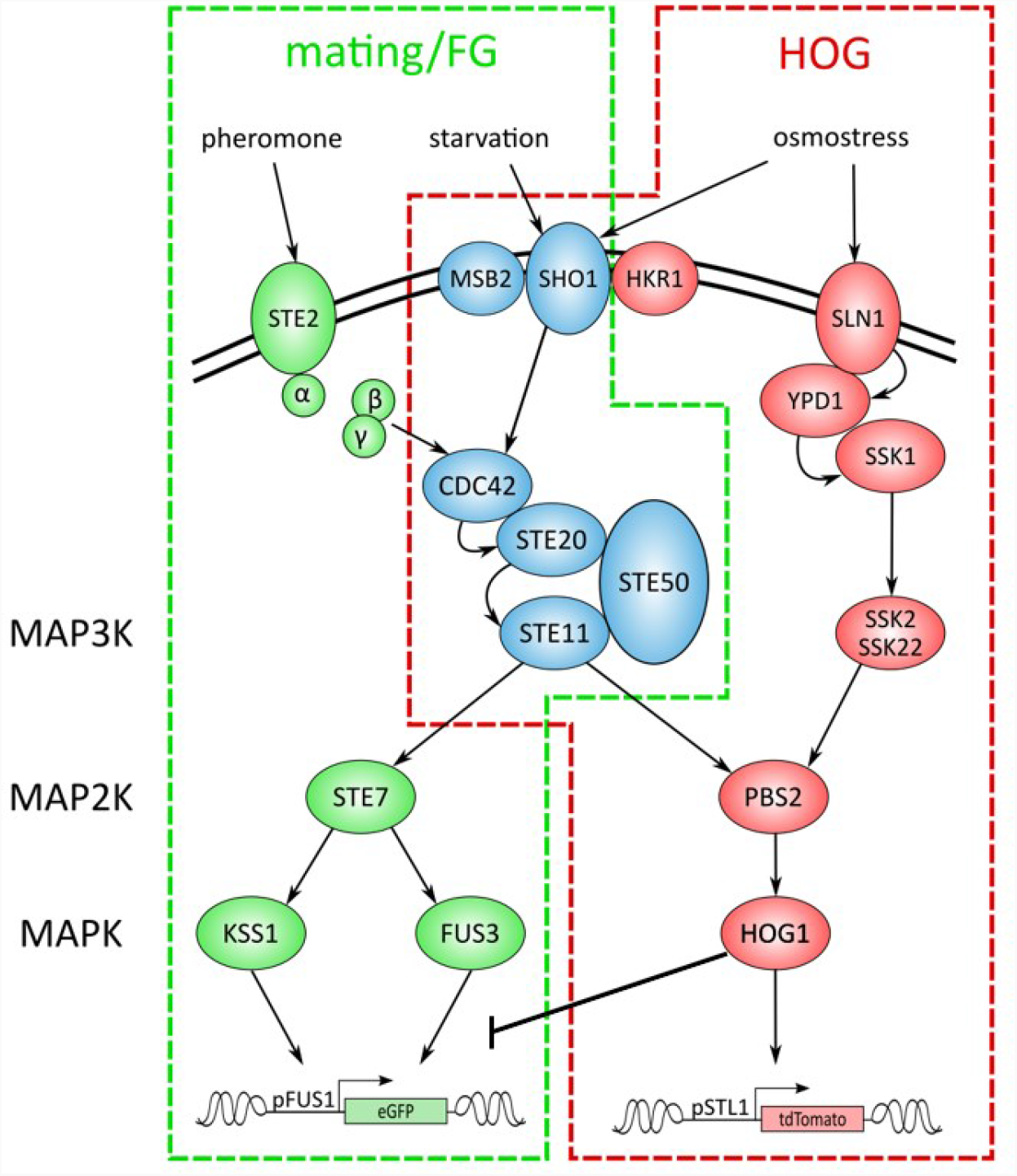
Yeast MAPK pathways. In yeast, the responses to pheromone, nutrient starvation, and hyperosmotic stress are controlled by mitogen activated protein kinase (MAPK) networks termed the mating pathway, filamentous growth (FG) pathway, and the high osmolarity glycerol (HOG) pathway respectively. The HOG pathway is outlined in red, and the mating/FG pathways are outlined in green. Components belonging to the mating pathway only are shown in green; components belonging to the HOG pathway only are shown in red; blue components belong to more than one of the three pathways. In our experiments, activation of the HOG pathway results in expression of pSTL1-tdTomato and mating/FG pathway activity results in expression of pFUS1-eGFP.

A striking feature of these networks is that they share an activation mechanism, namely the activation of Ste11p by Cdc42-Ste20 (Figure 1). Despite this connection, the pathways are (in general) insulated and it is thought that a combination of scaffolding and inhibition via Hog1p suppresses activation of the mating/FG pathways in response to osmostress [29–31]. This discovery initiated extensive research into how the connection between the two pathways is regulated, and various approaches have been used to study this phenomenon, the majority of which have involved large changes to the network structure. For example, it has been shown that deleting Hog1p results in significant leakage of osmostress signal into the mating/FG pathways (a phenomenon known as crosstalk) [29], as does chemical inhibition of Hog1p or rendering Hog1p kinase dead [32,33]. Similarly, crosstalk is seen when the MAP2K binding domains of Hog1p and Kss1p or Fus3p are swapped, effectively physically rewiring the networks [34]. How much, though, does signal insulation and crosstalk vary between wild-type strains, where there is natural genetic variation but the networks remain intact?

Here we quantitatively examine signaling in two strain backgrounds, YPH499 and Σ1278b (Sigma), and compare their response to osmostress and pheromone. YPH499 is congenic with S288C and has previously been used to study crosstalk between these pathways [35,36]. Sigma is commonly used to study filamentation and has long been used to study the mating and FG pathways [11,37,38]. These two strains are genetically very similar, diverging by ∼0.3% which is similar to the divergence between unrelated humans[39–41]. Whole genome sequencing studies have shown that while both strains are domesticated laboratory strains, Sigma is more significantly diverged from S288C [39,41]. Importantly, the components of the HOG and mating/FG MAPK pathways are present in both strains and therefore the two strains have the same apparent network structure.

We show that, despite the two strains having the same network structure, they show qualitative and quantitative differences in the response to osmostress and pheromone. This is true when the strains are stimulated with osmostress alone, and when the cells are exposed to simultaneous osmostress and pheromone. We also show that the well-known suppression of the mating/FG pathways by the HOG pathway activity can be separated into two phases, a previously unknown early, strain-dependent phase and the known late, Hog1p-dependent phase. In the early phase, crosstalk occurs in Sigma but not in YPH499 cells. We also show that a known HOG-dependent suppressor of crosstalk, kinase Rck2p, has a more significant role in Sigma than in YPH499. By comparing two strains, we demonstrate that different strains vary significantly in their signaling output, despite apparently identical network structures.

## Results

### Sigma and YPH499 have different osmosensitivity

We wished to quantitatively compare the effects of simultaneous osmostress and pheromone in the YPH499 and Sigma backgrounds. However, the effects of simultaneous induction are known to correlate with the severity of osmostress, and we therefore needed to establish the relative osmosensitivity of the two strains. When grown in liquid culture and on agar plates, Sigma is more osmosensitive than YPH499, though not grossly and both strains recover well from an osmotic shock.

When YPH499 and Sigma cells were spotted onto YPD agar with and without sorbitol, both strains showed decreased growth with increasing sorbitol (Figure 2A). Comparing the two strains after 24 hours growth, however, showed that Sigma grows comparatively worse than YPH499 at higher concentrations of sorbitol, with the defect clearly visible beginning at 0.75M sorbitol. This defect lessened after 45 hours growth, consistent with adaptation to the hyperosmolar environment. Quantification of spot growth confirms these general trends when the Sigma growth is normalized to YPH499 growth (Figure 2B). At 24 hours and at 45 hours, the relative growth of Sigma decreased linearly with increasing sorbitol indicating that more osmostress imposed a more severe growth defect in Sigma than in YPH499. At 24 hours, the slope of the best fit line is -0.41 M^-1^, indicating the growth defect on 1M sorbitol YPD agar was approximately 41% greater in Sigma than in YPH499. In contrast, the slope of the line at 45 hours is -0.19 M^-1^, meaning that the growth defect per molar sorbitol at 45 hours was only 19% greater than in YPH499.

**Figure 2:**
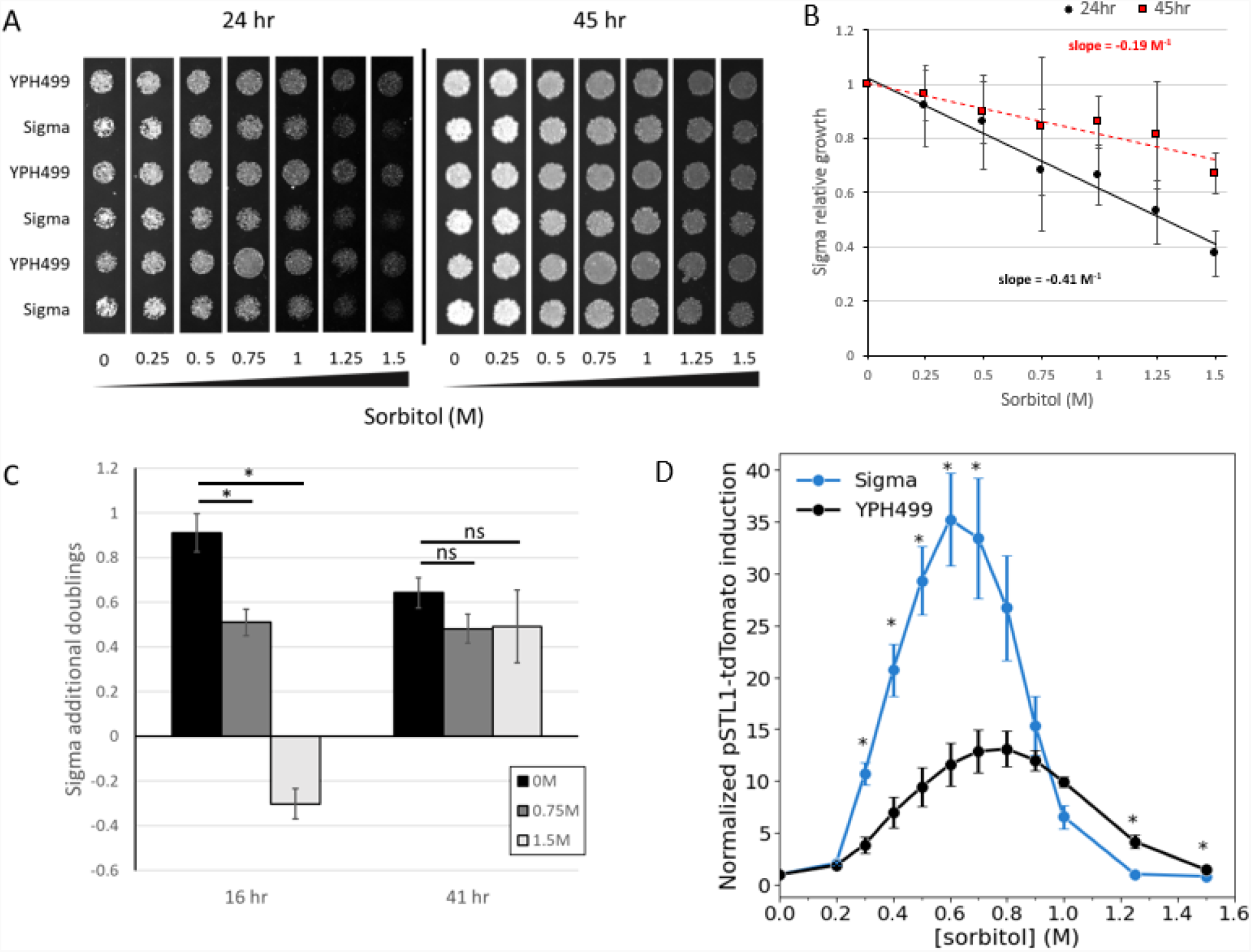
Sigma is more osmosensitive than YPH499. **A**: Spot tests of YPH499 and Sigma on varying concentrations of sorbitol after 24 hr (left) and 45 hr (right) at 30°C. **B**: Quantification of the growth defect in Sigma relative to YPH499 as a function of sorbitol concentration. The quantification was done at 24 hr (black circles) and at 45 hr (red squares). The slope of the line represents the relative growth defect in Sigma per molar sorbitol. Plotted are mean ± standard error of the mean (s.e.m.) of 3 biological replicates. **C**: Growth of YPH499 and Sigma in liquid YPD with or without sorbitol. OD600 measurements were used to calculate the number of doublings at 16 hr and 45 hr of 3 cultures each of Sigma and YPH499 in YPD with the indicated dose of sorbitol. Bars represent the mean ± s.e.m. of the difference between the Sigma doublings and the YPH499 doublings. p-values calculated using two-sided student’s t-test with equal variance. *p<0.05. n.s., not significant. **D**: Induction of pSTL1-tdTomato in YPH499 (gray) and Sigma (blue) after 45 minutes in various doses of sorbitol. Plotted are mean ± s.e.m. of 3 biological replicates.

Similar results were found in liquid culture (Figure 2C). We have consistently seen that, in several media types, Sigma yeast grows to a higher density than YPH499 yeast in liquid culture without osmostress (data not shown). To test the effects of osmostress in liquid growth, we measured optical density at 600 nm (OD600) after 16 hours growth and 41 hours growth in liquid YPD with and without sorbitol. After 16 hours growth in YPD without sorbitol, Sigma cultures were about twice as dense as YPH499 cultures (0.9 additional doublings). The addition of sorbitol to the media however significantly decreased Sigma’s growth advantage at 16 hours: at 0.75M sorbitol, the excess growth was reduced to 0.5 additional doublings, and at 1.5M sorbitol the Sigma cultures were *less* grown than the YPH499 cultures, having doubled 0.3 times fewer than the YPH499 cultures. After 45 hours, the effects of sorbitol are largely abrogated: the number of additional doublings in the cultures with sorbitol was slightly lower than the cultures without sorbitol although the difference was not statistically significant.

Finally, Sigma shows greater induction of a HOG pathway transcriptional reporter than YPH499 at most doses of sorbitol (Figure 2D). We measured induction of tdTomato driven by the *STL1* promoter (pSTL1-tdTomato) after 45 minutes in varying concentrations of sorbitol using flow cytometry. In both strain backgrounds maximum pSTL1-tdTomato induction was achieved at 0.7M – 0.8M sorbitol before decreasing at higher concentrations of sorbitol. In the Sigma background, however, maximum pSTL1-tdTomato induction was nearly 30-fold, while the maximum induction in YPH499 was only 15-fold. Additionally, pSTL1-tdTomato induction at high concentrations of sorbitol decreased more sharply in Sigma than in YPH499, and in particular pSTL1-tdTomato was less induced in Sigma at 1M and 1.25M sorbitol.

### Mating pathway activation under osmostress is faster in Sigma than in YPH499

Having seen that Sigma is more osmosensitive than YPH499, we reasoned that the effect of osmostress on mating pathway induction should be more severe in Sigma than in YPH499 at a given concentration of sorbitol. Increasing osmostress has been shown to both dampen and delay mating pathway activation when cells are simultaneously exposed to an osmolyte (such as sorbitol or NaCl) [42], and we therefore hypothesized that Sigma should show greater dampening and a longer delay in mating pathway activation than YPH499. To test this, we induced Sigma and YPH499 cells with 10 µM α-factor and varying concentrations of sorbitol (0M – 1.5M) and measured expression of eGFP driven by the *FUS1* promoter (pFUS1-eGFP). The time courses of mating response in the pheromone/sorbitol induced cells, plotted as a percentage of the pFUS1-eGFP level in a time-matched sample induced with pheromone alone, had a characteristic shape (Figure 3A). There is a large initial dampening in the mating response which gradually recovers to a less extreme final dampening level. We quantify the initial dampening by taking the minimum value of the mating response and the final dampening by taking the average of the final two measurements at 150 minutes and 180 minutes post-induction, by which point the cells had fully recovered from the osmotic shock. To quantify the delay, we find the full-width half minimum (FWHM) by calculating the width of the initial dampening peak at the dampening level halfway between the initial dampening and the final dampening. A greater FWHM indicates a longer time to recover to the final dampening level.

**Figure 3:**
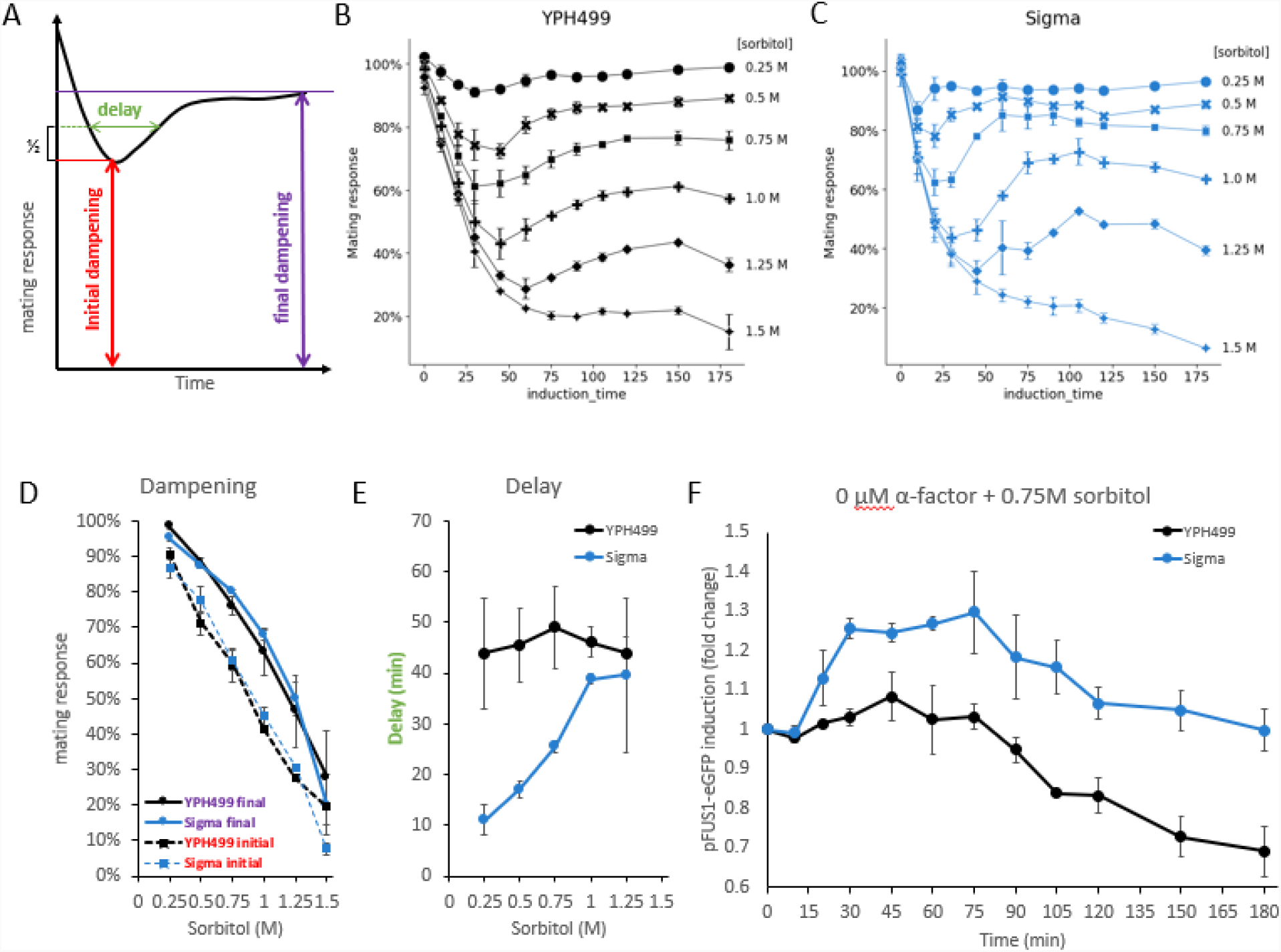
Effect of osmostress on mating pathway activation. **A**: Sorbitol disruption of the mating pathway consists of a strong initial dampening followed by recovery to a final dampening level. Initial dampening – minimum value of curve; final dampening – average of 150 min. and 180 min. time points. Delay is calculated as the full-width half minimum. **B**: Effect of various concentrations of sorbitol on mating pathway induction in YPH499 at 10 μM α-factor. Points and error bars are mean ± standard error of the mean (s.e.m.) of 3 biological replicates (0.25M – 0.75M) or 2 biological replicates (1M – 1.5M). **C**: Effect of various concentrations of sorbitol on mating pathway induction in Sigma at 10 μM α-factor. Points and error bars are mean ± s.e.m. of 3 biological replicates (0.25M – 0.75M) or 2 biological replicates (1M – 1.5M). **D**: Initial and final dampening due to osmostress are identical in YPH499 and Sigma. Points and error bars are mean ± s.e.m. of 3 biological replicates (0.25M – 0.75M) or 2 biological replicates (1M – 1.5M). **E**: Delay as a function of sorbitol in Sigma and YPH499. Sigma has a smaller delay at low-to-moderate concentrations of sorbitol. Points and error bars are mean ± s.e.m. of 3 biological replicates (0.25M – 0.75M) or 2 biological replicates (1M – 1.5M). **F**: Induction of pFUS1-eGFP in Sigma in response to sorbitol alone. Points and error bars are mean ± s.e.m. of 2 biological replicates.

In YPH499 (Figure 3B) and in Sigma (Figure 3C), increasing sorbitol caused increasing dampening in mating pathway activation, both initially and finally. Notably, we observed no strain-dependent difference in dampening (Figure 3D). This was true when considering initial dampening (dashed line) and final dampening (solid line) at every level of osmostress. In both strains, the minimum mating response (initial dampening) at 0.25M sorbitol was approximately 90% of a sample induced with pheromone alone. These samples recovered to a final mating response greater than 95% of the pheromone only sample (YPH499, 98.4%; Sigma, 95.1%). Under more severe osmostress (1.25M sorbitol), the initial dampening was much greater, approximately 30% in both strains (YPH499, 26.7%; Sigma, 30.6%). Similarly, the final dampening was also more severe, and the cells recovered to only 50% of a pheromone only sample (YPH499, 46.4%; Sigma, 50.1%). At the most severe osmostress tested (1.5M sorbitol), the cells did not recover to a final dampening level but remained depressed for the duration of the time course. The final dampening level at this osmostress is 27.8% in YPH499 and 20.0% in Sigma.

Unlike the dampening effect, the delay in mating activation caused by osmostress was clearly strain-dependent (Figure 3E). In YPH499, the delay of the mating response curves was roughly constant at 45-50 minutes at each test sorbitol concentration. In contrast, the delay in Sigma increased with increasing sorbitol, from 11 minutes at 0.25M sorbitol to 40 minutes at 1.25M sorbitol. We were unable to calculate the FWHM for our 1.5M sorbitol time courses because the curves did not recover to a final dampening level but instead decreased throughout the time course.

### Sigma yeast show induction of the mating pathway under osmostress

The finding that Sigma yeast have a starkly shorter delay in mating pathway activation under simultaneous osmostress and pheromone induction was surprising because it is known that activation of the Hog1p inhibits induction of mating pathway transcripts [29,32,33,36,43]. Counter to this, in examining a sorbitol-only control, we noticed that Sigma increased pFUS1-eGFP expression under sorbitol induction, even in the absence of pheromone (Figure 3F). When Sigma and YPH499 cells were exposed to 0.75M sorbitol, the Sigma cells showed a maximum of 1.3-fold increase in pFUS1-eGFP at approximately 30-45 minutes post-induction. In contrast, the maximum induction in YPH499 was negligible (1.05-fold), and its induction was uniformly lower than the induction seen in Sigma. This finding was unexpected, and we performed further experiments to verify that Sigma is showing mating pathway induction under osmostress. We collected time courses in a variety of mating and HOG pathway mutants following a 0.75M sorbitol induction (the results of which are detailed below) in 24-well plates which are less susceptible to edge effects and well-to-well variability than the 96-well plates used in the simultaneous sorbitol and pheromone experiments. These experiments were performed in an identical manner, and the wild-type controls represent 20 biological replicates performed on different days. The collected controls from these mutant experiments (Figure 4A) show that Sigma induced pFUS1-eGFP under osmostress significantly more than YPH499. Sigma achieved maximum induction at approximately 45 minutes post-induction; at this time point, the pFUS1-eGFP induction in Sigma cells (normalized to the 0-minute time point) was 1.24-fold compared to 1.08-fold in YPH99, and pFUS1-eGFP induction in Sigma was significantly higher than in YPH499 at every time point from 20 minutes to 60 minutes post-induction. We also measured pFUS1-eGFP induction at different concentrations of sorbitol at 45 minutes post-induction (Figure 4B). pFUS1-eGFP induction in Sigma was strongly dose-dependent, achieving its maximum induction at a moderate level of osmostress, approximately 0.6M-0.8M sorbitol. In contrast, YPH499 cells did not show significant pFUS1-eGFP induction at any concentration of sorbitol. As seen in the pSTL1-tdTomato dose-response curve in Figure 2D, the pFUS1-eGFP induction in Sigma decreased at higher concentrations of sorbitol and we saw negligible induction at concentrations greater than 1M sorbitol.

**Figure 4:**
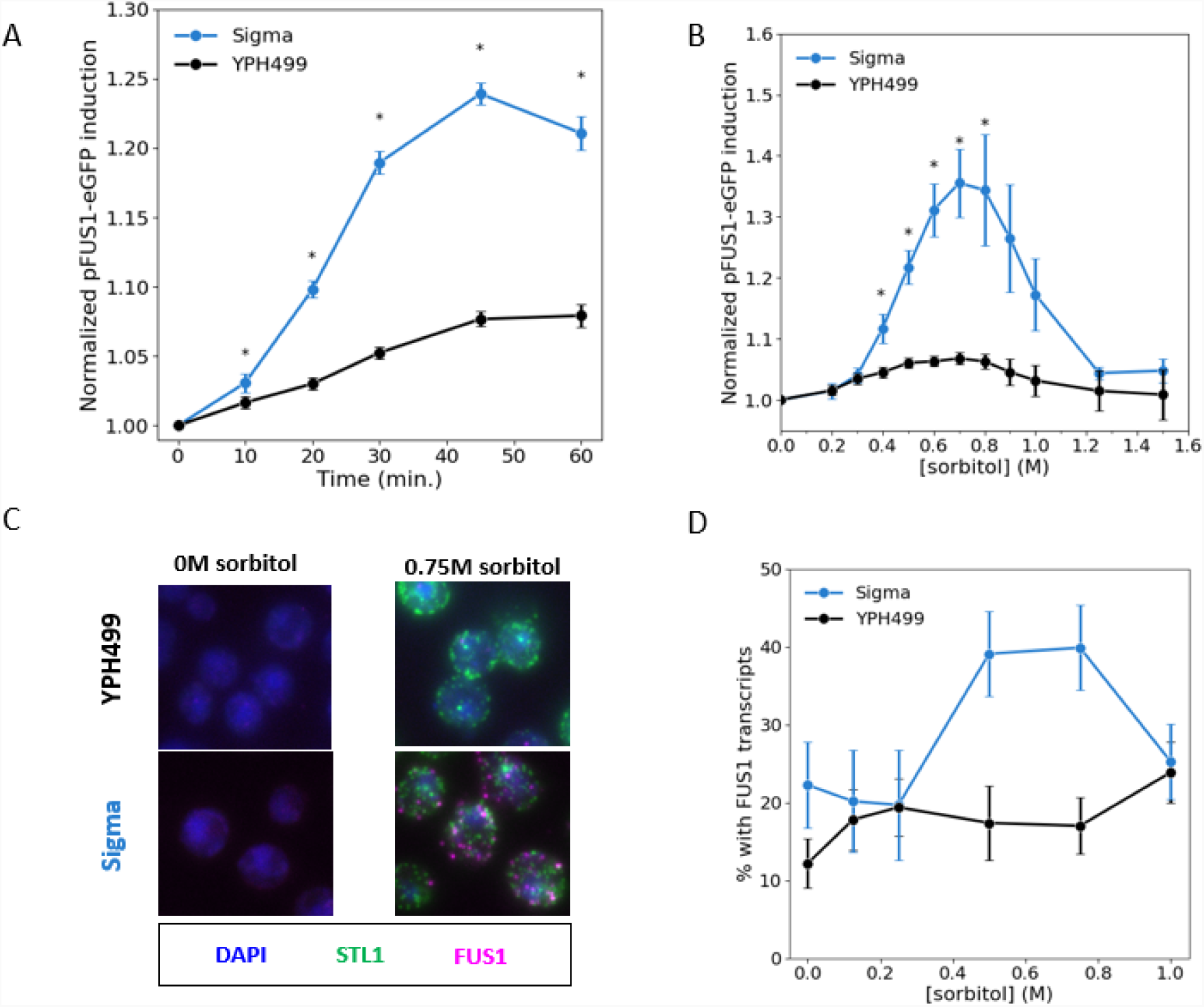
Sigma induces the mating pathway at certain levels of osmostress. **A**: Flow cytometry time course of sorbitol-induced pFUS1-eGFP induction in YPH499 and Sigma. Data for each strain is normalized to the t = 0 time point. (mean +/- s.e.m.; N = 19 biological replicates; Sigma t=60 is 17 biological replicates). * p<0.05, two-sided student’s t-test with equal variance. **B**: pFUS1-eGFP induction at 45 min. in the indicated concentration of sorbitol. Data for each strain is normalized to the [sorbitol] = 0 point. (mean +/- s.d.; n = 3 biological replicates). * p<0.05, two-sided student’s t-test with equal variance. **C**: Representative images of cells after a 10 minute induction with 0M sorbitol (left) or 0.75M sorbitol (right). Single mRNA transcripts appear as green (STL1) or magenta (FUS1) dots. Cells were additionally stained with DAPI (blue). **D**: Proportion of cells with at least 1 FUS1 transcripts after 10 min induction with varying doses of sorbitol. Error bars represent at 95% confidence interval for the proportion of cells with FUS1 transcripts. Average N = 313 cells in each FISH experiment.

To confirm that this apparent induction was happening at the level of transcription, we directly observed *FUS1* transcripts using single-cell fluorescent *in situ* hybridization (scFISH). In both strain backgrounds, there were few *FUS1* transcripts present in an uninduced sample (Figure 4C, left images). In Sigma, however, many cells had FUS1 transcripts following a 15-minute induction with 0.75M sorbitol (Figure 4C, bottom right). We did not see the same in YPH499 cells (Figure 4C, top right). As seen with flow cytometry, induction of *FUS1* transcripts under sorbitol in Sigma was dose dependent and at 0.5M and 0.75M sorbitol, roughly 40% of cells had *FUS1* transcripts, an increase from 20% at 0M sorbitol (Figure 4D). In YPH499 however, no dose-dependent induction was observed and the number of cells with *FUS1* transcripts remained at 10-20% at every concentration of sorbitol.

### Sorbitol induction of pFUS1-eGFP in Sigma is *STE11* and *FUS3/KSS1* dependent

We wondered whether the pFUS1-eGFP induction in Sigma represented true crosstalk, this is, whether the induction was due to signal leaking from the HOG pathway into the mating/FG pathways. To test this, we collected sorbitol induction time courses in mating/FG pathway deletions. The Sigma-background deletions are discussed here. Ste11p is a shared component of the HOG and mating pathways and can phosphorylate either Pbs2p (in the HOG pathway) or Ste7p (in the mating/FG pathways). Deleting *STE11*, therefore, should disrupt the connection between the pathways and prevent pFUS1-eGFP induction due to osmostress in the Sigma background. As expected, pFUS1-eGFP induction was drastically reduced in Sigma *ste11Δ* cells (Figure 5A). In fact, in a 60-minute time course under 0.75M sorbitol, pFUS1-eGFP induction in Sigma *ste11Δ* cells was indistinguishable from that in YPH499 wild-type cells. Activation of Ste11p may plausibly result in phosphorylation of either Fus3p or Kss1p, and activation of either kinase would cause an increase in pFUS1-eGFP production. We therefore collected time courses in *fus3Δ* and *kss1Δ* deletions as well as a *fus3Δ kss1Δ* double deletion. As seen in the *ste11Δ* deletion, pFUS1-eGFP induction in Sigma *fus3Δ kss1Δ* cells was reduced to levels seen in YPH499 wild-type cells (Figure 5B). This indicates that Fus3p and/or Kss1p activity is responsible for the increased pFUS1-eGFP induction seen in the Sigma background under osmostress. In contrast, we found that pFUS1-eGFP induction in the Sigma background was unchanged when *FUS3* and *KSS1* were deleted separately (Figure 5C, D). This is true for the duration of the 60-minute time course. This suggests that signal leaks from Ste11p into both kinases and activation of either kinase is sufficient to induce our reporter.

**Figure 5:**
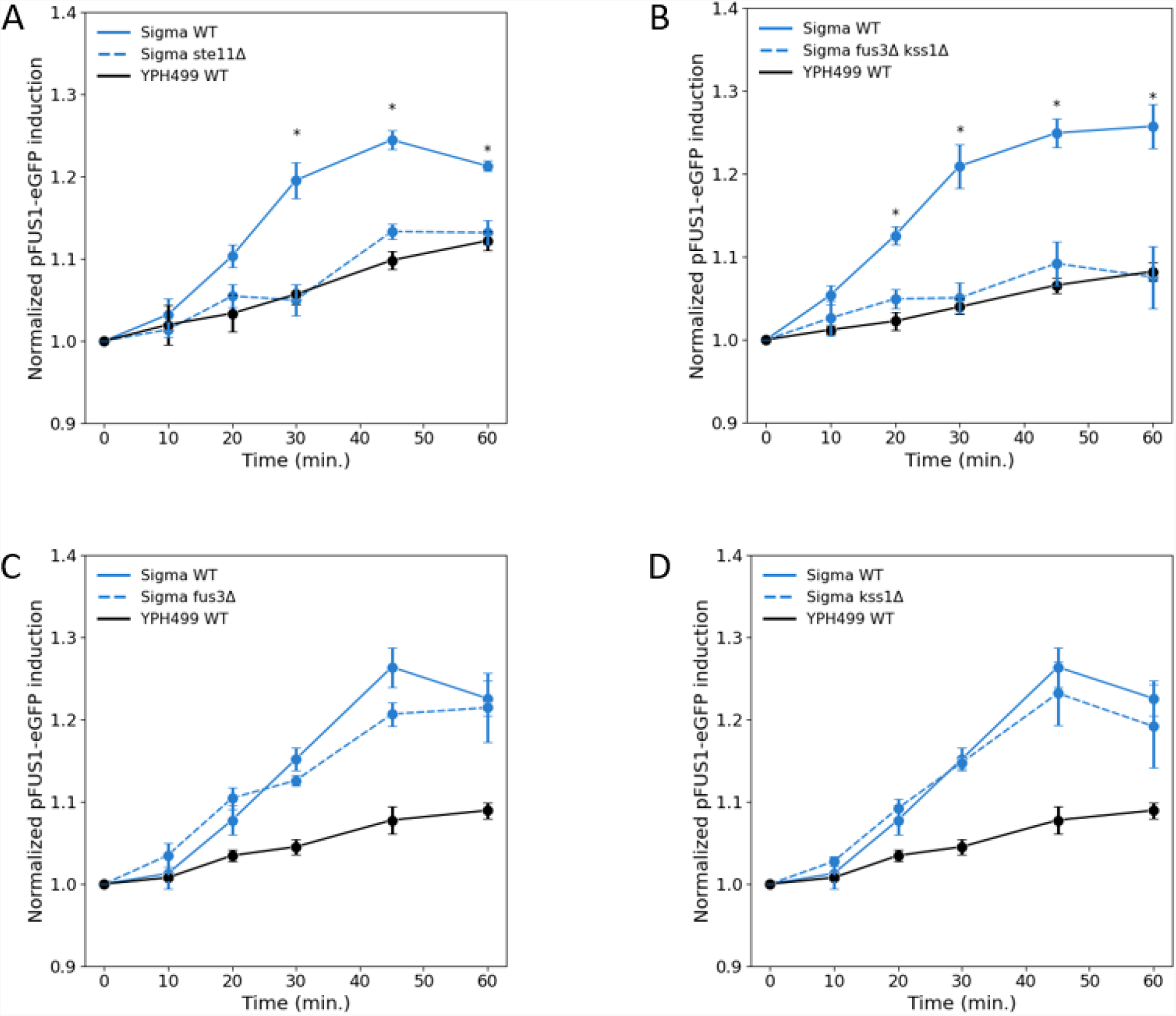
Crosstalk in Sigma is dependent on mating pathway components. Flow cytometry measurements of crosstalk in mating/FG pathway deletions. Each panel shows a Sigma background deletion (**A**: fus3Δ; **B**: kss1Δ; **C**: ste11Δ; **D**: fus3Δ kss1Δ) along with Sigma-background and YPH499-background wild-type controls. Measurements are normalized to the t = 0 timepoint. Plotted are mean +/- s.e.m.; n = 3 (panels A, C, D) or 4 (panel B) biological replicates. The wild-type controls were performed concurrently with the deletion in each panel. The fus3Δ and kss1Δ deletions (panels C and D) were measured simultaneously and the wild-type control lines are the same in both panels. *p<0.05, student’s t-test with equal variance.

### Mechanisms of HOG-dependent crosstalk inhibition are strain dependent

HOG pathway activity is known to suppress crosstalk in part through activation of the MAPK activated protein kinase Rck2p, perhaps through translational suppression of mating pathway components [42]. We wondered whether a defect in Rck2p activation could explain the crosstalk seen in the Sigma background. We collected 60-minute time courses under 0.75M sorbitol in YPH499 and Sigma wild-type, *rck2Δ*, and *hog1Δ* cells (Figure 6). In both strains, the *rck2Δ* and the *hog1Δ* deletions each produced excess crosstalk compared to the wild-type cells. Interestingly, this excess crosstalk occurred late in the induction time course, and, in particular, it occurred after the crosstalk seen in the Sigma wild-type cells. In the YPH499 background, both deletions showed a significant increase in pFUS1-eGFP induction compared to the wild-type beginning at 30 minutes post-induction (Figure 6A). In the Sigma background, the *rck2Δ* showed excess induction beginning at 30 minutes post-induction and the *hog1Δ* showed additional crosstalk beginning at 45 minutes post-induction (Figure 6B). We also found that the role of *rck2Δ* in the YPH499 background was limited compared to the Sigma background. Specifically, the maximum amount of crosstalk in the YPH499 *rckΔ* was considerably lower than in the YPH499 *hog1Δ* cells (*rck2Δ* = 1.32 ± 0.05; *hog1Δ* = 2.00 ± 0.04). In contrast, Sigma *rck2Δ* cells achieved roughly the same level of crosstalk as the *hog1Δ* cells over 60 minutes (*rck2Δ* = 1.66 ± 0.07; *hog1Δ* = 1.69 ± 0.08), though the signal was clearly attenuating in the *rck2Δ* mutant.

**Figure 6:**
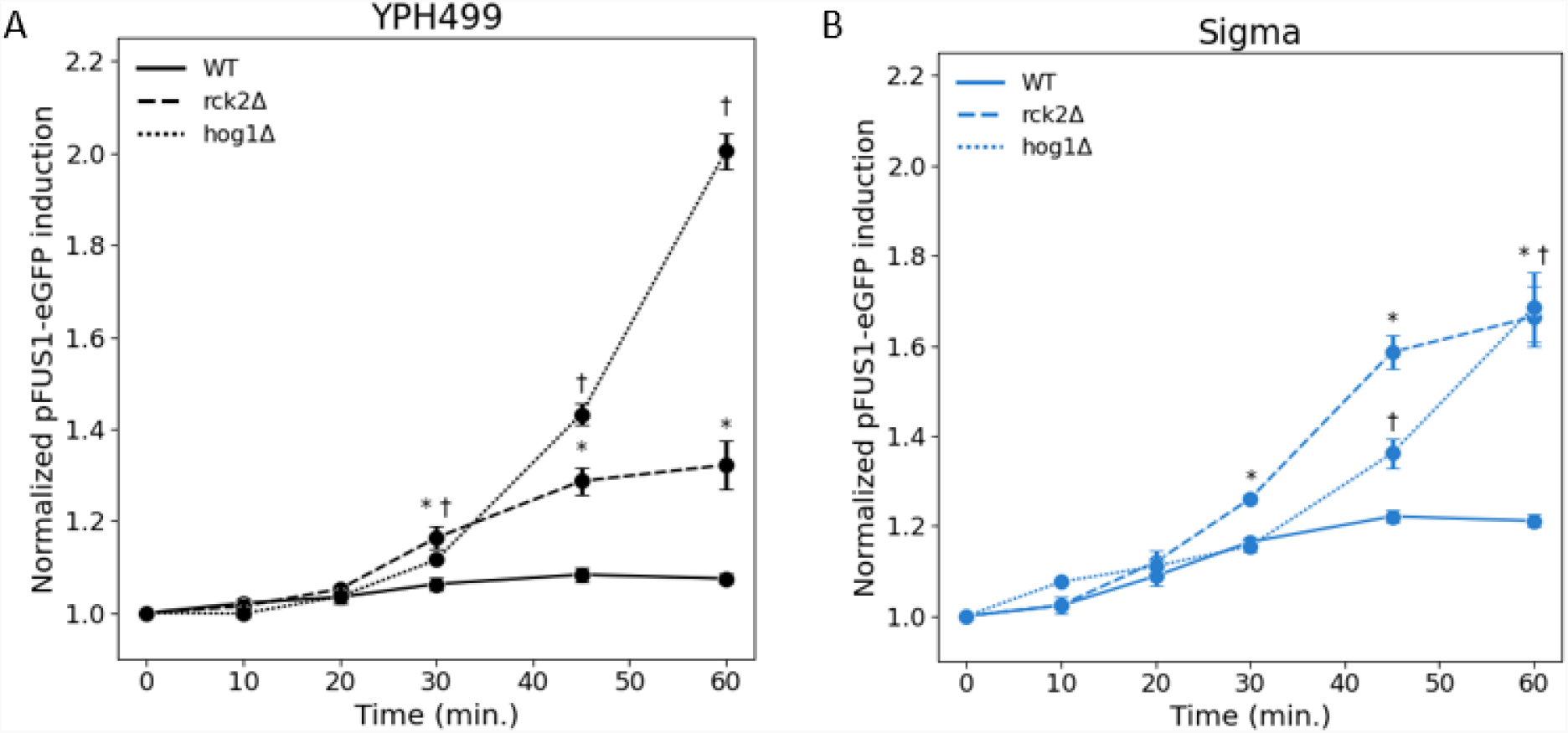
HOG-dependent suppression of crosstalk occurs late in a time course. Flow cytometry measurements of crosstalk in WT (–), *hog1Δ* (…), or *rck2Δ* (--) cells in the (A) YPH499 or (B) Sigma backgrounds. Measurements are normalized to the t = 0 time point. Mean ± s.e.m.; n = 3 biological replicates. A student’s t-test with equal variance was performed between WT and *rck2Δ* (*) and between WT and *hog1Δ* (†). Markers represent points at which p<0.05.

### Early crosstalk in Sigma is independent of HOG pathway activity

Having seen that deleting *HOG1* produces crosstalk late in an induction time course, we wondered whether other disruptions of Hog1p activity also produce crosstalk late in a time course. We measured crosstalk in two additional mutants in which Hog1p activity is disrupted. First, we examined *ssk1Δ* deletions. Ssk1p is an essential component of the fast-activating SLN1 branch of the HOG pathway and deleting *SSK1* significantly slows activation of Hog1p [44–46]. In the Sigma background, the *ssk1Δ* cells and wild-type cells showed identical pFUS1-eGFP induction for the first 30 minutes post-induction, but the *ssk1Δ* cells showed increased induction at 45 minutes and 60 minutes post-osmotic shock (Figure 7A). In the YPH499 background, results were similar. The *ssk1Δ* cells showed higher induction than wild-type cells at 45- and 60-minutes post-induction (Figure 7C). There was also a slight but significant increase in induction at 30 minutes post-shock in the YPH499 background that we did not observe in the Sigma background.

**Figure 7:**
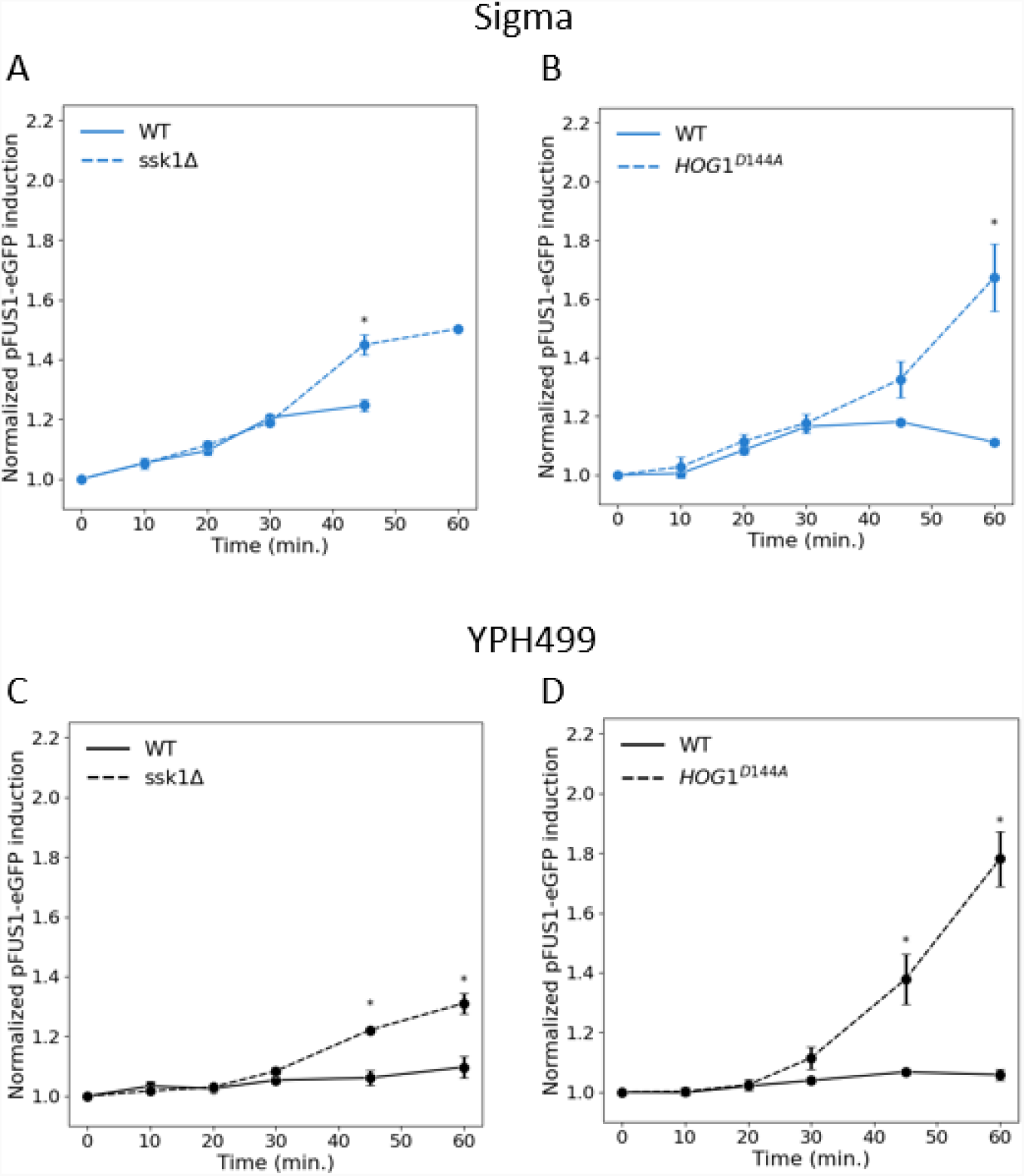
HOG pathway disruptions affect late crosstalk but not early crosstalk. Flow cytometry measurements of crosstalk in mutants which disrupt HOG pathway activity. Each panel shows a Sigma background or YPH499 background deletion of *SSK1* (panels **A, C**) and a kinase-dead *HOG1*^*D144A*^ mutant (panels **B, D**) along with a wild-type control. Measurements are normalized to the t = 0 time point. The wild-type controls were performed concurrently with the deletion in each panel. Mean +/- s.e.m.; N = 3 biological replicates. *p<0.05, student’s t-test with equal variance.

We also disrupted Hog1p activity directly by introducing the kinase-dead *HOG1*^*D144A*^ allele which displays no kinase activity [32]. In the Sigma background the *HOG1*^*D144A*^ mutant cells and wild-type cells exhibited identical levels of pFUS1-eGFP induction for the first 30 minutes following the osmotic shock, but the mutants showed higher crosstalk at 45 and 60 minutes (Figure 7B). Similarly, in the YPH499 background the *HOG1*^*D144A*^ mutants showed much higher induction at 45 minutes and 60 minutes post-induction with a slight increase at 30 minutes post-induction (Figure 7D). In both strain backgrounds, and similarly to the *hog1Δ* time courses, pFUS1-eGFP induction in the *HOG1*^*D144A*^ mutants increases continuously throughout the time course while the induction is limited in the *ssk1Δ* cells. Because the HOG pathway disruptions do not affect the crosstalk seen in Sigma during the first 30 minutes of an induction, we conclude that the mechanism which permits early crosstalk in the Sigma background is not dependent on HOG pathway activity.

## Discussion

### Osmostress has strain dependent effects on mating pathway activation

The two strains used in this study, Sigma and YPH499, show differences in osmosensitivity. Sigma grows worse than YPH499 on YPD agar and in liquid YPD when sorbitol is present, and Sigma shows stronger activation of a HOG pathway transcriptional reporter than YPH499 at most concentrations of sorbitol. This result is not unexpected because Sigma contains functional aquaporins Aqy1p and Aqy2p while these genes in YPH499 contain a premature stop codon rendering them less functional than the Sigma variant [47–49]. Consequently, Sigma cells are more prone to water loss following an osmotic shock. The loss of Aqy1p and Aqy2p function has been directly tied to increased osmotolerance in growth on YPD agar and in hyperosmotic/hypoosmotic cycling [47,48]. It has been shown that the water loss at high concentrations of sorbitol significantly slows diffusion of signaling molecules and slows HOG pathway activation [50]. This is the likely cause of the decreased pSTL1-tdTomato induction at high concentrations of sorbitol (Figure 2D). Sigma is predicted to lose water more readily than YPH499 due to its functional aquaporins, which would explain both the increased transcriptional activation at lower concentrations of sorbitol (because the stress is more severe) and the decreased activation at high concentrations of sorbitol (because slowed diffusion is more severe).

As has been previously reported, simultaneous osmostress and pheromone results in a significant dampening and delay to mating pathway activation [42]. This is expected because activation of the HOG pathway is fast relative to mating pathway activation and Hog1p activity strongly inhibits the mating pathway. Because Hog1p activity (both amplitude and duration) correlates with osmostress [45], the simplest explanation of the dampening and delay effect is that it is a direct result of Hog1p activity under osmostress. Surprisingly, we find that the dampening effect is identical in our two strain backgrounds, despite Sigma being more osmosensitive. This is true at every tested concentration of sorbitol, ranging from a small osmostress to a severe osmostress, and is true both initially and at saturation. Despite the similarity in dampening, our results indicate that the delay is strongly strain dependent. Sigma has a shorter delay at every concentration of sorbitol, and the delay in Sigma is dose dependent while it is constant in YPH499. Nagiec and Dohlman found a dose dependent delay in the S288C background (with which YPH499 is congenic), so it is curious that we observe a constant delay in our experiments [42]. A potential explanation for this is that our strains harbor a *bar1Δ* deletion while the strains used by Nagiec do not. Bar1p is a protease which degrades pheromone [51,52] and its expression is induced by mating pathway activation as a form of negative feedback [53,54]. Consequently, the pheromone activation time courses saturate after a few hours in *BAR1* strains but do not in *bar1Δ* strains. Thus, our measure of delay is based on the time it takes doubly stimulated cells to produce pFUS1-eGFP at the same rate as cells induced with pheromone alone, instead of being a direct measure of time-to-saturation as measured by Nagiec and Dohlman. Nonetheless, the difference in delay seen in the two strains is striking and indicates that inhibition of the mating pathway in Sigma is not directly related to the severity of osmostress as has been previously suggested.

### HOG pathway signal transiently leaks into the mating pathway in Sigma yeast

We find transient induction of the mating/FG pathways under osmostress alone in Sigma yeast, and this induction is consistent with signal leakage from the HOG pathway into the mating/FG pathways through the Ste11p node. This was unexpected because the known targets of Hog1p activity which suppress the mating pathway, namely Ste50p and Rck2p, are present in Sigma and the structure of the HOG and mating pathways is the same in both strains. This result indicates that the insulation between the HOG and the mating/FG pathways is strain dependent. Crosstalk is typically studied in the context of a large disruption to the signaling networks (for example, deletion of Hog1p or deletion of Hog1p targets), largely because crosstalk is not observed in wild-type cells. Here we show that crosstalk can occur in a native network absent large disruptions, but it is strain-dependent and transient. Importantly, this crosstalk occurs under physiologically relevant stress profiles. For example, during wine-making yeast must survive an initial sugar content of 15-28% (w/v) [55], which assuming glucose is the primary sugar, produces an osmoshock equivalent to 0.8M - 1.6M sorbitol. A previous study observed crosstalk into a mating/FG reporter under complex, quickly oscillating stress [56]. Here, our stress is a simple, static shock which more likely to be found in a natural environment. Further study of this phenomenon will provide insight into the factors which contribute to or suppress crosstalk in a fully functioning network.

### Rck2p-dependent inhibition of crosstalk is strain dependent

Rck2p is an important factor in HOG-dependent suppression of crosstalk. The exact mechanism by which Rck2p suppresses crosstalk is unknown, but it has been suggested that activation of Rck2p translationally suppresses activation of mating pathway products and disrupts the positive feedback necessary to fully activate the mating pathway [42]. We found that in Sigma, deleting *RCK2* produces similar levels of crosstalk as deleting *HOG1*. In fact, crosstalk in the *rck2Δ* mutant was higher than in the *hog1Δ* at some time points. This suggests that Rck2p is the factor responsible for suppressing late crosstalk in the Sigma background. In contrast, we observed in YPH499 that crosstalk is attenuated relative to *hog1Δ* cells, and while *rck2Δ* cells produce excess crosstalk relative to WT cells, this crosstalk is significantly lower than the crosstalk seen in *hog1Δ* cells. A previous study (in the S288C background) showed that Rck2p is not solely responsible for HOG-dependent suppression of crosstalk [42], so it is surprising that crosstalk in Sigma *rck2Δ* cells reaches similar levels to that seen in Sigma *hog1Δ* cells. Further experiments are needed to understand the differences in the strain backgrounds which cause this difference in Rck2p-mediated insulation.

### The HOG pathway regulates late, but not early, crosstalk

Our initial hypothesis for the crosstalk seen in the Sigma background was that a defect in HOG pathway signaling allowed some transient crosstalk prior to robust activation of Hog1p. Our results, however, suggest that, while HOG pathway activity inhibits late crosstalk, it does not explain the early crosstalk seen in the Sigma background. In the *ssk1Δ* mutants, the fast-activating SLN1 branch of the HOG pathway is disrupted, and it has been shown that activation of the HOG pathway is roughly two-fold slower in *ssk1Δ* cells [44–46]. The *hog1Δ* and *HOG1*^*D144A*^ mutants both eliminate Hog1p activity, the former by eliminating Hog1p itself and the latter by rendering Hog1p kinase dead [32]. We used both mutations because the MAPK pathway kinases are promiscuous and will phosphorylate any target that can bind [34,57,58], therefore removing Hog1p as a binding partner for Pbs2p may impact signaling through the network for reasons other than Hog1p activity. If the crosstalk in Sigma were due to ineffective Hog1p activity, we expect to see excess crosstalk in the mutants at the time points at which we see crosstalk in Sigma. In all three cases, we *do* see additional crosstalk, but it occurs *after* the crosstalk seen in wild-type cells. That is, the time courses of *ssk1Δ, hog1Δ*, and *HOG1*^*D144A*^ mutants are identical to wild-type time courses for 30 minutes post-induction but show increased induction at the 45- and 60-minute time points. In YPH499, these mutants show slightly increased crosstalk at 30 minutes and substantially increased crosstalk at 45- and 60-minutes post-induction.

This result allows us to divide crosstalk into two phases: early and late. Early following an osmotic shock (roughly the first 30 minutes), crosstalk is permitted in the Sigma background but not the YPH499 background. Late in the time course, beginning at roughly 30 minutes, additional crosstalk is suppressed by Hog1p activity in both strain backgrounds. Because Hog1p activity is merely delayed and not eliminated in *ssk1Δ* cells, our findings suggest that there is a critical window following osmostress in which Hog1p must be active in order to suppress late crosstalk. Without a functioning SLN1 branch, activation of Hog1p occurs outside this window and late crosstalk is permitted, though it is attenuated as the cell responds to the stress and turns off the HOG pathway. This model agrees with a previous study which found that Hog1p needed to be active for 20 to 30 minutes in order to suppress crosstalk, but that inhibition of Hog1p after 30 minutes did not cause crosstalk, indicating that continuous Hog1p activity is not needed to suppress crosstalk throughout an osmotic shock [36].

The factors which permit early crosstalk in Sigma but suppress early crosstalk in YPH499 are unknown and further studies are necessary to determine how these factors are regulated. Our results indicate that suppression via these factors is not Hog1p dependent because crosstalk in the HOG-disrupted mutants remained low at 20- and 30-minutes post-induction in the YPH499 background and was unchanged in the Sigma background. One potential HOG-independent mechanisms of crosstalk inhibition is scaffolding [30]. However, there is only a single point mutation in Ste5p (the mating pathway scaffold [59] in the Sigma background (D877G) which occurs outside any kinase binding sites; Pbs2p (the HOG pathway scaffold [7,60]) is identical in the two strains. Further, our results show that Kss1p is sufficient to allow crosstalk in the Sigma background, and activation of Kss1p does not require a scaffold [61]. Thus, differences in the scaffold proteins themselves are unlikely to be the cause of the early crosstalk seen in the Sigma background, though further studies are needed to conclusively eliminate this mechanism.

## Conclusion

Studies of signaling in yeast frequently use mutations or deletions to discover how different nodes in a signaling network affect signal flow through the network. While these experiments yield invaluable information about network structure and function, they also introduce large changes into the network. For example, deleting *HOG1* not only eliminates Hog1p activity but also eliminates Pbs2p’s binding partner and target and significantly impairs the cell’s ability to adapt to osmostress. We were interested in what differences in signaling can exist in cells which retain the native network structure. We have quantitatively compared two strains which ostensibly have identical HOG and mating pathway structure; that is, neither strain harbors a deletion of any MAPK component. Despite this, we found differences in MAPK signaling between these strains. The Sigma background is, compared to the S288C-congenic YPH499 background, more osmosensitive when measured through growth on solid YPD agar, growth in liquid YPD, and expression of an osmostress-responsive reporter. The increased osmosensitivity, though, does not cause increased dampening of the mating pathway when cells are costimulated with osmostress and pheromone, and, surprisingly, the Sigma background has a shorter delay in mating pathway activation at most concentrations of sorbitol. We also observe crosstalk from the HOG pathway into the mating pathway when wild-type Sigma cells are stimulated with osmostress alone, a result which to our knowledge has not been reported in any *S. cerevisiae* strain. Our results indicate that this signal leakage occurs through Ste11p and Fus3p/Kss1p, which is consistent with the known links between the HOG and mating pathways. Importantly, this crosstalk occurs early in an osmostress induction time course, prior to the known HOG-dependent inhibition of crosstalk. Finally, we have also shown that known mechanisms of crosstalk inhibition may vary differ between strain backgrounds. The crosstalk seen in Sigma background *rck2Δ* cells is comparable to that seen in Sigma *hog1Δ* cells, while in the YPH499 background *rck2Δ* exhibit significantly lower crosstalk than *hog1Δ* cells. Our results demonstrate that a careful comparison of laboratory yeast strains can provide insight into how signaling is regulated in the context of a natural, undisrupted signaling network. Undoubtedly there is some difference in the Sigma and YPH499 networks which permits crosstalk, but this difference could be the result of a subtle mechanism, such as differences in expression or point mutations caused by natural genetic variation, and not the result of a large disruption to the network. Future work to discover how other strains regulate the HOG and mating pathways, particularly in comparison to S288C-derived strains, will provide insight into the diverse mechanisms by which signaling may be regulated.

## Materials and Methods

### Yeast strains and methods

Yeast culture and growth were performed using standard methods [62] and transformations were done using a lithium acetate transformation protocol [63]. For scFISH and flow cytometry experiments, cultures were grown in low fluorescence media (1.7 g/L Yeast Nitrogen Base without ammonium sulfate, without folic acid, without riboflavin [MP Biomedicals # 114030512]; 5 g/L ammonium sulfate; 20 g/L dextrose) supplemented with amino acids. Other experiments were performed in standard YPD media or synthetic complete media with the appropriate amino acid dropped out.

Genes were deleted using homologous integration of a drug selection cassette amplified from pMM0129, pMM0130, or pMM0131 and verified by colony PCR. CRISPR/Cas9-mediated allele replacements were performed by first deleting the gene of interest with a drug selection cassette followed by transformation of a CRISPR plasmid expressing a guide RNA targeting the drug resistance gene and repair template amplified from a plasmid or genomic DNA as appropriate [64] (CRISPR plasmids were a gift from Audrey Gasch). Transformant colonies were passaged 3X in YPD media to lose the CRISPR plasmids and the integration of the allele was confirmed by sequencing. Where necessary, drug selection markers on plasmids were exchanged using circular polymerase extension cloning [65,66]. A complete list of strains and plasmids can be found in Tables S1 and S2.

The YPH499-background strain with integrated pSTL1-tdTomato and pFUS1-eGFP fluorescent reporters (yMM0736) was a gift from Jeremy Thorner, as were the plasmids containing these reporters (pMM0154 and pMM1055). The pFUS1-eGFP reporter was amplified from pMM0154 and cloned into the pYIPlac211 backbone between the BamHI and EcoRI sites. Sigma-background strain with these reporters were constructed as described [36] by digesting the reporter plasmids with an enzyme which cuts once in the promoter (NruI for *STL1* and BsaAI for *FUS1*) prior to integration. This method integrates the reporter alongside the original (undisrupted) gene.

To facilitate experiments in 96-well plates, the Sigma background was rendered non-clumpy by introducing the *AMN1*^*D368V*^, which allows daughter cells to cleanly separate from mother cells [67–69] (Figure 8). *AMN1*^*D368V*^ was introduced into yMM1174 using CRISPR/Cas9 as described above.

**Figure 8:**
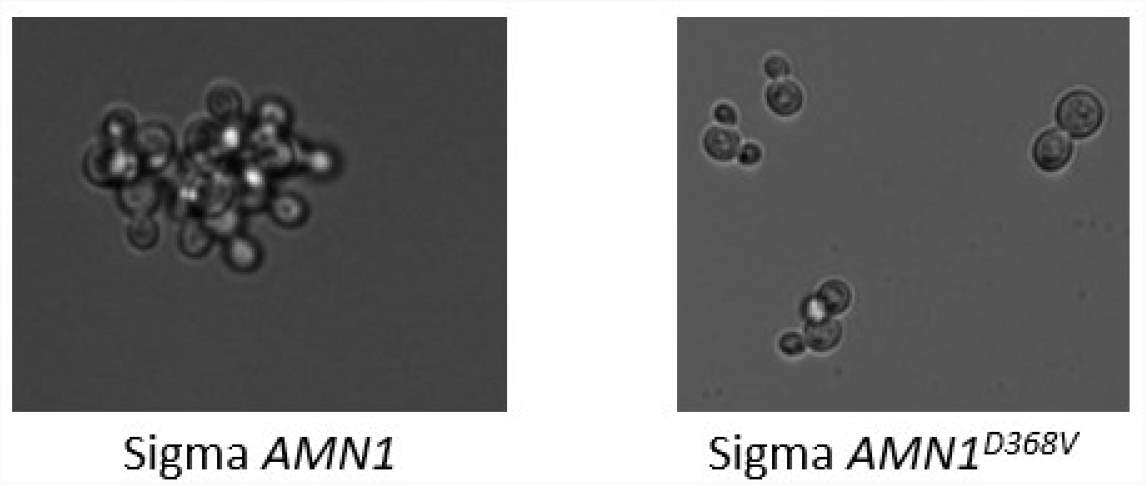
*AMN1* affects clumpiness in Sigma. Introducing the *AMN1*^*D368V*^ allele significantly reduces clumpiness in the Sigma background.

### Osmosensitivity

To quantify osmosensitivity on YPD agar, three colonies each of YPH499 and Sigma were grown to mid-log and spotted YPD plates with various concentrations of sorbitol. Plates were imaged using a ChemiDoc (Bio-Rad) at 24 hr and 45 hr and spot intensity was quantified using the gel analyzer tool in ImageJ. Intensity on YPD+sorbitol plates was normalized to intensity on the YPD (no osmostress) plate to determine growth, and the resulting mean growth of the Sigma spots was divided by the mean growth of the YPH499 spots to determine the growth defect of Sigma relative to YPH499. The standard error of the mean was found for the growth quantification and was propagated in the relative growth defect calculating using the standard formula.

To quantify osmosensitivity in a well-mixed culture, 3 colonies each of YPH499 and Sigma were grown to mid-log then diluted to approximately 1.5 × 10^6^ cells/mL in YPD with or without sorbitol. Cultures were incubated with rotation at 30°C and samples were taken at 16 hr and at 41 hr. Density was quantified as the optical density at 600 nm using a spectrophotometer and samples were diluted to within the linear range of our spectrophotometer prior to measurement. Doublings were calculated by taking 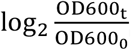, and the number of doublings in YPH499 was subtracted from the number of doublings in Sigma to calculate the excess doublings in Sigma. The standard error of the mean for the doublings was found and propagated in the excess doublings calculation using the standard formula.

### Induction time courses

Overnight cultures of the appropriate yeast strains were diluted to 1.5-3 × 10^6^ cells/mL in LFM and allowed to grow 6-8 hr at 30°C to the mid-log phase. To induce, a sample of the mid-log culture was pipetted into a 96-well (Figure 3) or 24-well plate (Figure 2D, 4A,B, 5-8) containing media with sorbitol and/or pheromone. After inducing for the desired time, cycloheximide was added to a final concentration of 100 μg/mL to halt translation. Plates were sealed with BreatheEasy film and incubated shaking at 30°C for 16-18 hr to allow fluorescent proteins to mature. For time courses, samples were induced beginning with the latest timepoint and continuing in reverse so that cycloheximide was added to all samples at the same time and all samples were allowed to fold for the same amount of time. Cycloheximide was added to the well containing the 0-minute time point prior to induction. For experiments in 24-well plates (using clumpy yMM1174-derived Sigma strains), 3X volume PBS+0.1% Tween-20 (pre-chilled to 4°C) was added to each well and plates were placed on ice. each well was sonicated (amplitude 5, 1 sec on, 1 sec off, 1 min total) before 225 μL was transferred to a 96-well plate pre-loaded with 25 µL 0.1% methylene blue as a live-dead stain. For experiments in 96-well plates (using non-clumpy yMM1584-derived Sigma strains), 4X volume PBS+0.1% Tween+0.0125% methylene blue was added to each well and plates were immediately placed on ice. Samples were kept on ice and in the dark prior to flow cytometry.

### Flow cytometry

Samples were run on an Attune NxT Focusing Cytometer. Measurements were taken in the forward scatter (FSC), side scatter (SSC), BL1 (488nm laser, 530/30 filter, 503LP dichroic mirror), YL1 (561nm laser, 585/16 filter, 577LP dichroic mirror) and RL2 (633 nm laser, 720/30 filter, 690LP dichroic mirror) channels. Samples were first gated for eGFP+ (BL1H) and tdTomato+ (YL1H) events which removed a significant portion of the debris and dead cells. Samples were then gated for cells using FSC-A vs SSC-A, then two successive gates for single cells were drawn using FSC-A vs FSC-H and SSC-A vs SSC-H respectively. Finally, a gate for live cells was drawn using RL2-H vs FSC-H to gate for low RL2 (live) cells. See supplementary figure 9 for a representative example of the gating procedure in YPH499 (Figure 9A) and Sigma (Figure 9B). At least 50,000 total events were collected for each sample. The median BL1-H and YL1-H were taken as the eGFP value and the tdTomato value respectively for each sample. For double stimulus experiments, eGFP values of the sorbitol + pheromone induced samples were normalized to a time-matched sample stimulated with pheromone alone. For crosstalk inductions, the eGFP values were normalized to a time-matched control sample induced with 0M sorbitol media to account for differences in basal pFUS1-eGFP production. All analysis was performed using FlowJo version 10.8.

**Figure 9:**
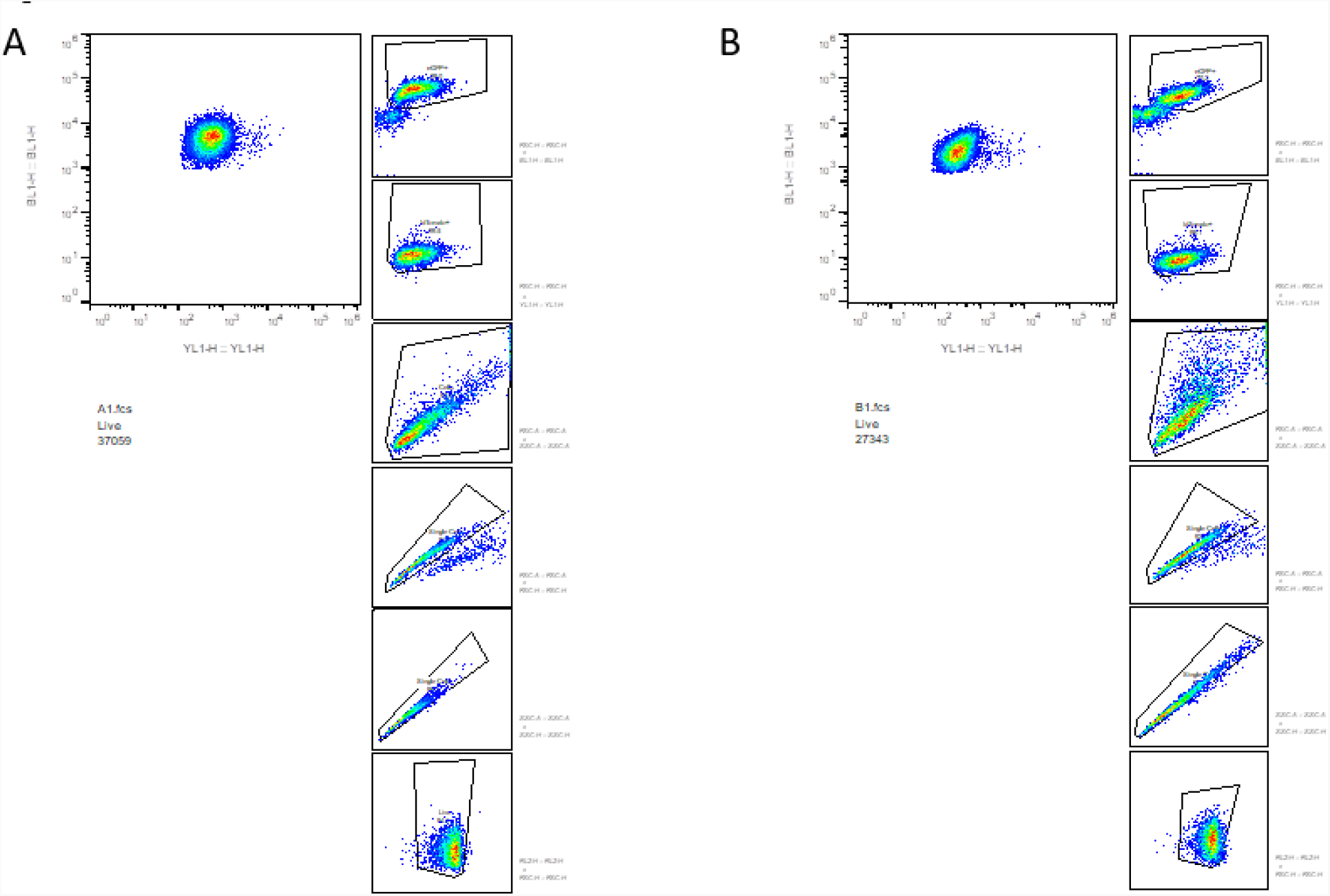
Flow cytometry gating. Representative examples of flow cytometry gating in the (**A**) YPH499 and (**B**) Sigma backgrounds.

### Fluorescent *in situ* hybridization

Strains were grown as described above and collected for fixation and processing as described in [70]. FISH probes were designed using the Biosearch Technologies Stellaris Designer with Quasar 670 (*FUS1* probes) or Quasar 570 (*STL1* probes). Probes sequences are available in tables S3 and S4. Images were acquired as z-stacks every 0.2 mm with an epifluorescent Nikon Eclipse-TI inverted microscope using a 100x Nikon Plan Apo oil immersion objective and Clara CCD camera (Andor DR328G, South Windsor, Connecticut, United States of America). Quasar 670 emission was visualized at 700 nm upon excitation at 620 nm (Chroma 49006_Nikon ET-Cy5 filter cube, Chroma Technologies, Bellows Falls, Vermont, USA). Quasar 570 emission was visualized at 605 nm upon excitation at 545 nm (Chroma 49004_Nikon ET-Cy3 filter cube). Transcripts were counted by semiautomated transcript detection and counting in MATLAB using scripts adapted from [71].

