## Supporting Information for "Strain dependent differences in coordination of yeast signaling networks"

### Table S1: List of yeast strains

| Strain | Background | Genotype | Source |
|---|---|---|---|
| yMM0736 (YJP212) | YPH499[1] | MATa pSTL1::HA-tdTomato::ADE2 pFUS1::HA-eGFP::ADE1 bar1Δ::KanMX | [1] |
| yMM1052 (YJP406) | YPH499 | yMM0736 ssk1Δ::TRP1 | [1] |
| yMM1053 (YJP407) | YPH499 | yMM0736 sho1Δ::HygMX | [1] |
| yMM1062 | YPH499 | yMM0736 fus3Δ::HygMX | This study |
| yMM1109 | YPH499 | yMM0736 ste11Δ::HygMX | This study |
| yMM1159 | YPH499 | yMM0736 kss1Δ::HygMX | This study |
| yMM1160 | YPH499 | yMM0736 fus3Δ::HygMX kss1Δ::NatMX | This study |
| yMM1722 | YPH499 | yMM0736 hog1Δ::HygMX | This study |
| yMM1723 | YPH499 | yMM0736 HOG1(D144A) | This study |
| yMM1724 | YPH499 | yMM0736 rck2Δ::HygMX | This study |
| yMM0964 | Sigma[2] | MATa | David Botstein |
| yMM0985 | Sigma | MATa bar1Δ::NatMX | This study |
| yMM1124 | Sigma | MATa leu2Δ0 ura3Δ0 | David Botstein |
| yMM1127 | Sigma | MATa leu2Δ0 ura3Δ0 pSTL1::HA-tdTomato::LEU2 | This study |
| yMM1128 | Sigma | MATa leu2Δ0 ura3Δ0 pSTL1::HA-tdTomato::LEU2 bar1Δ::NatMX | This study |
| yMM1174 | Sigma | MATa pSTL1::HA-tdtomato::LEU2 pFUS1::HA-eGFP::URA3 bar1Δ::NatMX | This study |
| yMM1179 | Sigma | yMM1174 sho1Δ::HygMX | This study |
| yMM1180 | Sigma | yMM1174 ste11Δ::HygMX | This study |
| yMM1181 | Sigma | yMM1174 ssk1Δ::HygMX | This study |
| yMM1182 | Sigma | yMM1174 kss1Δ::HygMX | This study |
| yMM1183 | Sigma | yMM1174 fus3Δ::HygMX | This study |
| yMM1185 | Sigma | yMM1174 fus3Δ::HygMX kss1Δ::KanMX | This study |
| yMM1583 | Sigma | yMM1174 amn1Δ::KanMX | This study |
| yMM1584 | Sigma | yMM1174 AMN1(D368V) | This study |
| yMM1725 | Sigma | yMM1174 hog1Δ::HygMX | This study |
| yMM1726 | Sigma | yMM1174 HOG1(D144A) | This study |
| yMM1727 | Sigma | yMM1174 rck2Δ::HygMX | This study |

[1] YPH499 genotype: MATa ura3-52 lys2-801_amber ade2-101_ochre trp1-Δ63 his3-Δ200 leu2-Δ1 [1]

[2] Sigma2000, MATa prototroph derived from Σ1278b [2]

### Table S2: List of plasmids

| Plasmid | Description | Source |
|---|---|---|
| pMM0004 | pRS406 | [1] |
| pMM0025 | pRS316 HOG1 | Sharad Ramanathan |
| pMM0129 | pFA6 NatMX | [3] |
| pMM0130 | pFA6 HygMX | [3] |
| pMM0131 | pFA6 KanMX | [3] |
| pMM0154 | YIPlac128 pFUS1-HA-eGFP | [4] |
| pMM0155 | YIPlac128 pSTL1-HA-tdtomato | [4] |
| pMM0292 | YIPlac211 | [5] |
| pMM0300 | pYIPlac211 pFUS1-HA-eGFP | This study |
| pMM0887 | pXIPHOS-NatMX pSNR52-KanMX sgRNA-tSNR52 | Audrey Gasch |
| pMM0888 | pXIPHOS-NatMX pSNR52-HygMX sgRNA-tSNR53 | Audrey Gasch |
| pMM0889 | pXIPHOS-NatMX pSNR52-tSNR54 | Audrey Gasch |
| pMM0890 | pXIPHOS-HygMX pSNR52-KanMX sgRNA-tSNR55 | This study |
| pMM0891 | pXIPHOS-HygMX pSNR52-tSNR56 | This study |
| pMM0904 | pHS2-KanMX | This study |
| pMM1232 | pXIPHOS-KanMX pSNR52-HygMX sgRNA-tSNR53 | This study |
| pMM1233 | pXIPHOS-KanMX pSNR52-tSNR54 | This study |
| pMM1234 | pRS316 HOG1$^{D144A}$ | This study |

### Table S3: *STL1* FISH probe sequences

| Sequence | Name |
|---|---|
| cagtgactggttctgcttat | STL1_1 |
| caacttcttacccgtaagtc | STL1_2 |
| cgtcatagatgcgatagtga | STL1_3 |
| tcgtatccaaacagggagaa | STL1_4 |
| tagacttgccatcaaccctt | STL1_5 |
| ccattttctttggttgctgg | STL1_6 |
| agttgcgtgtctgtcatgat | STL1_7 |
| tcataacaggaggttgtagc | STL1_8 |
| agaacctgcgaaacaaccta | STL1_9 |
| caccgcagaacataacgaat | STL1_10 |
| gaacccatcaggattaatgg | STL1_11 |
| cggcaccaatgatggttatt | STL1_12 |
| cacgaaatgcgcatgtagaa | STL1_13 |
| ataaactggcctaatgccca | STL1_14 |
| attcaaccctgttccaacac | STL1_15 |
| gccaaacgggaatagtagat | STL1_16 |
| gcaaccctctattttcagct | STL1_17 |
| ccaaaagcaattgtggaacc | STL1_18 |
| ctgttggtataagacaaccc | STL1_19 |
| tttgcattgacacggggaat | STL1_20 |
| agcaggaagagagcaaaaac | STL1_21 |
| gtggcgattcaggtagttta | STL1_22 |
| cgactttgagaaatcagcca | STL1_23 |
| taccaagtagcgagcttctt | STL1_24 |
| ttcctcatcatttggatccg | STL1_25 |
| gtgaagcatagcaacttctg | STL1_26 |
| gtttggtcctgttaacagca | STL1_27 |
| ggagaacaaacttgacagtg | STL1_28 |
| tctgaagattttgggacctg | STL1_29 |
| gttgaagctgcaatcaaagc | STL1_30 |
| gcagcgttacaaccagtaaa | STL1_31 |
| ccccacctatgatcattgat | STL1_32 |
| aaggcgtagattgttgcgaa | STL1_33 |
| gcttacgtctacctagcttt | STL1_34 |
| gacctgtggcacctaataaa | STL1_35 |
| gcacctcttgcgttttcttt | STL1_36 |
| gaacaaaaataagccgacgg | STL1_37 |
| ggtgggtatatccatggtaa | STL1_38 |
| gcacgaactttcattgatgc | STL1_39 |
| tgtggagaaagcgtttgttg | STL1_40 |
| ccgcaaagttacacaaccaa | STL1_41 |
| caaccggactgtccaataaa | STL1_42 |
| cggtttcagggtagaaaaag | STL1_43 |
| gtcgatttcctccaaacttc | STL1_44 |
| cctcgtatgctttagcaaag | STL1_45 |
| tagcaactctccatggttga | STL1_46 |
| agggataacttgggcaaatg | STL1_47 |
| ggcatgatcttcgacttctt | STL1_48 |

### Table S4: *FUS1* FISH probe sequences

| Sequence | Name |
|---|---|
| gtcgtctgcattattgttgc | FUS1_1 |
| taaggtagtagacattgcgg | FUS1_2 |
| ggaactagcttgcgaagata | FUS1_3 |
| cgttactgttgttacactcg | FUS1_4 |
| gagcgtattgatgtcgctat | FUS1_5 |
| ccgccacattagaaaagagt | FUS1_6 |
| gctgaagatgattttggctg | FUS1_7 |
| gattgaaagcccaattgtgc | FUS1_8 |
| cagaatattccgatgggaag | FUS1_9 |
| gaaatggacaccgaattcct | FUS1_10 |
| ttggaatcgtagctgacatg | FUS1_11 |
| ttagtgcggcgacaatattc | FUS1_12 |
| ccaaaataaccgtgagaacc | FUS1_13 |
| tctgatcctcacacttactc | FUS1_14 |
| gggtgtcgttatacttctca | FUS1_15 |
| aagacatgttatcacccgag | FUS1_16 |
| gcggtattatgggtttggaa | FUS1_17 |
| ccgttttcttaggtgtcagt | FUS1_18 |
| gaccaagcatatgggttctt | FUS1_19 |
| ctttggggtctaacgaaatg | FUS1_20 |
| ctttctcttcctccatttcg | FUS1_21 |
| gtatacaggaatgcatccac | FUS1_22 |
| gcatgctggattcaatatgg | FUS1_23 |
| gacaccgtttttgagaagg | FUS1_24 |
| ctcactcgtttttaacgcag | FUS1_25 |
| gtggagattcgtaactccat | FUS1_26 |
| tagaacccctcaagaaccat | FUS1_27 |
| ggggtctttaaagaggtctt | FUS1_28 |
| atttgcttcagttgtggagc | FUS1_29 |
| acttatccgggagagcattt | FUS1_30 |
| ctgactcattgaagtaaccc | FUS1_31 |
| gatcgttcgtcaggcattat | FUS1_32 |
| gaggcgtgttattatactcc | FUS1_33 |
| cccaagttattcacactgtc | FUS1_34 |
| ttgtgaatctggcgtggtat | FUS1_35 |
| gccgtgattagatcgatgtt | FUS1_36 |
| gtatatcactgaatggggtc | FUS1_37 |
| ctcgtgtttctcagtgcttt | FUS1_38 |
| ttgatggggtgggtattatc | FUS1_39 |
| gcaatggtttagaacgtgac | FUS1_40 |
| cctccccattatatttggag | FUS1_41 |
| atgtcttccctaattggacg | FUS1_42 |
| ggctcgtaatcctgaataac | FUS1_43 |
| ccagcgagattcttatttcg | FUS1_44 |
| ggtatgagtggccagaattt | FUS1_45 |
| tttctaccagacaccatcca | FUS1_46 |
| ctgacgtgaatagaaccctt | FUS1_47 |
| gcctctatcttcattgaggt | FUS1_48 |
